# Eye-movement artifact correction in infant EEG: A systematic comparison between ICA and Artifact Blocking

**DOI:** 10.1101/2024.03.19.585723

**Authors:** Diksha Srishyla, Sara Jane Webb, Mayada Elsabbagh, Christian O’Reilly, The BASIS Team

## Abstract

**Background:** Independent Component Analysis (ICA) is a well-established approach to clean EEG and remove the impact of signals of non-neural origin, such as those from muscular activity and eye movements. However, evidence suggests that ICA removes artifacts less effectively in infants than in adults. This study systematically compares ICA and Artifact Blocking (AB), an alternative approach proposed to improve eye-movement artifact correction in infant EEG.

**Methods:** We analyzed EEG collected from 50 infants between 6 and 18 months of age as part of the International Infant EEG Data Integration Platform (EEG-IP), a longitudinal multi-study dataset. EEG was recorded while infants sat on their caregivers’ laps and watched videos. We used ICA and AB to correct for eye-movement artifacts in the EEG and calculated the proportion of effectively corrected segments, signal-to-noise ratio (SNR), power-spectral density (PSD), and multiscale entropy (MSE) in manually selected EEG segments with and without eye-movement artifacts.

**Results:** On the one hand, the proportion of effectively corrected segments indicated that ICA corrected eye-movement artifacts (sensitivity) better than AB. SNR and PSD indicated that both AB and ICA correct eye-movement artifacts with equal sensitivity. MSE gave mixed results. On the other hand, AB caused less distortion to the clean segments (specificity) for SNR, PSD, and MSE.

**Conclusions:** Our results suggest that ICA is more sensitive (i.e., it better removes artifacts) but less specific (it distorts clean signals) than AB for correcting eye-movement artifacts in infant EEG.

## 1. Introduction

EEG recordings contain high amounts of signals unrelated to cortical processing, such as muscular and ocular artifacts. This problem is particularly acute with recordings from infants, which are often not long enough for well-established approaches such as Independent Component Analysis (ICA) to work effectively [1]. Depending on the research question, a minimum recording length of 3.2 minutes has been suggested to apply ICA [2]. Yet, collecting *>* 3.2 minutes of usable EEG is often challenging in infants due to movement and fussiness. Further, although 3.2 minutes is a lower bound on the required length, the recommended length can be much longer. This recommended length can be determined with the formula *k · N* ^2^*/f*_*s*_, where *N* is the number of channels, *f*_*s*_ is the sampling frequency, and *k* is a multiplier whose value needs to be determined experimentally [2]. Using a test sample where ICA worked effectively, a value of 30 has been suggested for *k* [3]. In this example, the authors used EEG recorded from 32 channels, and they suggested that the appropriate value for *k* increases with the number of channels. Using the conservative *k* = 30 for a recording using 128 channels with a sampling rate of 500 Hz, at least 983 s would be required, an expectation often unrealistic for infant EEG recording. Further, ICA may not identify components related to eye movement in infant EEG [1].

Significant progress has been made in developing preprocessing pipelines addressing these issues. For example, the Maryland Analysis of Developmental EEG (MADE) pipeline removes residual eye-movement artifacts by first epoching the continuous data, then rejecting any epochs where the electrooculogram (EOG) channels exceed a certain voltage threshold [4]. Epochs are then examined and rejected if more than 10% of the EEG channels exceed a fixed voltage threshold. For the remaining epochs, EEG channels exceeding this threshold are interpolated.

The Lossless pipeline from the EEG Integrated Platform (EEG-IP-L) [5] applies a voltage-based criterion to identify periods where the data would not be corrected effectively by ICA due to non-stationary artifacts. It also tests for invalid channels based on extreme correlations between nearest neighbors and ensures a robust re-referencing on a consistent average reference regardless of the recording montage. It then applies multiple ICA decompositions, followed by an automated classification of components as neural or artifactual according to ICLabels – a classifier for independent components trained on adult data [6] – and an optional visual confirmation. This procedure for artifact removal is compatible with short recording durations, as is often the case with infant EEG, and can process epoched or continuous EEG alike [5]. This pipeline has recently been refactored and ported to Python to increase its usability and user-friendliness [7].

The Newborn EEG Artifact Removal (NEAR) [8] pipeline relies on Artifact Subspace Reconstruction (ASR), an approach designed originally for mobile adult EEG [9]. It detects invalid channels and adapts ASR to infant EEG by including a bad-channel detection, a necessary step for ASR effectiveness. ASR requires users to specify a parameter called ‘k’ and can either correct or remove artifacts. NEAR calibrates these parameters for effective artifact removal in infant EEG. This approach is more effective than ICA in removing artifacts with variable patterns of scalp topography and temporal dynamics [8]. The HAPPE (Harvard Automated Processing Pipeline for Electroencephalography) pipeline uses wavelet-thresholding ICA to mitigate distortions to spectral power estimates caused by ICA [2]. However, only a subset of channels can be used in high-density recordings (e.g., 128 channels) if the recording is not long enough for reliable ICA decomposition. As mentioned previously, that length can go up to more than 16 minutes for a sampling rate of 500 Hz recordings with 128 channels.

Also, efforts have been made to adapt the ICLabels classifier to infant EEG for automatic independent component labeling. The ADJUST classifier automatically identifies independent components by simultaneously calculating multiple features associated with the topographical and spectral properties of eye-movement artifacts [10]. However, this classifier is designed to work with event-related EEG paradigms, making it incompatible with continuous EEG processing. The Multiple Artifact Rejection Algorithm (iMARA) successfully classifies components into neural and artifactual categories when applied to both event-related and continuous (e.g., naturalistic) EEG recordings from infants [11].

As can be concluded from this overview, many pipelines rely on ICA for artifact identification and correction. As this approach has not been optimized for short infant EEG sessions, it is unclear to what extent ICA distorts the neural signals in EEG. A previous study suggested that Artifact Blocking (AB) distorts less clean EEG than ICA while effectively correcting for high amplitude artifacts [12]. However, this demonstration was qualitative and used only one EEG recording. In this study, we compare ICA with AB to quantitatively benchmark the performance of the two methods. We analyze resting state EEG recordings from 50 typically developing infants by manually annotating saccadic eye-movement artifacts, applying ICA and AB, and comparing the effectiveness of these algorithms based on i) visual inspection, ii) signal-to-noise ratio (SNR), iii) power spectral density (PSD), and iv) multiscale entropy (MSE) [12]. With this investigation, we aim to provide critical empirical evidence to help researchers decide which artifact correction method to use for future studies.

## 2. Methods

### 2.1. Participants

Recordings from the International Infant EEG Data Integration Platform (EEG-IP) were used for our analysis [5, 13]. EEG-IP includes EEG recordings from longitudinal infant-sibling studies conducted separately at two sites, London and Seattle [14, 15, 16, 17, 18]. EEG was recorded at ages 6-7 months and 12-14 months at both sites and 18 months in Seattle. Each site contributed raw EEG and behavioral data to this repository. Participants in these studies were grouped as having an elevated likelihood of Autism Spectrum Disorder (ASD) if they had an older sibling diagnosed with ASD or as comparison-controls with a typical likelihood of ASD if they did not. The repository comprises 420 recordings from 191 (94 females) infants.

For this study, we used recordings from the control group to benchmark AB and ICA on a sample representative of the overall infant population. We selected randomly 50 recordings for analysis. This sample was considered sufficient given that differences between methods tend to be reliable and systematic. Further, given this sample size, we can expect to detect 80% of the time with a 95% confidence an effect of 0.40 (Cohen’s d; paired t-test), which corresponds to an effect size small (0.2) to medium (0.5) [19]. We argue this is reasonable because very small effect sizes would mean that the difference between the approaches would be negligible in practice. Thus, we are mostly interested in effect sizes that are at least medium.

**Table 1:** Number of EEG recordings included in analysis by age, site and sex.

| Site | 6 months (male) | 12 months (male) | 18 months (male) |
| --- | --- | --- | --- |
| Seattle | 14 (11) | 16 (8) | 8 (6) |
| London | 12 (11) | 0 | 0 |

### 2.2. Recordings and pre-processing

EEG was recorded at a sampling frequency of 500 Hz with an EGI 128-channel Hydrocel net while infants sat on their caregivers’ laps and watched videos of brightly colored toys producing sounds or an adult woman singing nursery rhymes. An average reference was used.

Open-source software was used to preprocess, standardize the data, and minimize differences across studies/sites. Specifically, we adopted the Brain Imaging Data Structure (BIDS) extension to EEG [20, 21] and used the EEG-IP Lossless Pipeline to standardize the preprocessing [5]. This pipeline includes a semi-automatic preprocessing procedure to identify unreliable EEG signals and build comprehensive data quality annotations. It first addresses differences across datasets by executing a *staging script* specific to each project, including procedures for the co-registration of electrode coordinates to a common head surface, a robust average reference, and a 1Hz high-pass and notch filter (London: 49-51Hz; Seattle: 59-61Hz). The staging scripts then flag noisy periods and channels based on consistently outlying variance values to make files more comparable for later stages of the pipeline. Following the staging scripts, the pipeline assesses signal quality using confidence intervals of signal properties within each file to flag unusual periods and channels. Each time a problematic channel is flagged, the data are rereferenced using an average reference computed using channels interpolated on the shared co-registered head surface. The standard average reference ensures that differences in channel distributions (e.g., a 128-channel EGI net covers head regions not covered by less comprehensive montages) do not systematically impact the properties of the average reference. Following channel assessment, a robust Adaptive Mixture Independent Component Analysis (AMICA) procedure [22] is performed, and components are automatically classified as being or not of neural origin. The AMICA estimation used the EEGLAB plugin and default EEGLAB parameters. A single AMICA decomposition was used for independent component classification. The automatic classification relies on a crowd-sourced database (ICLabels) of labeled components, including 200,000 independent components from more than 6000 EEG recordings [6]. These components included artifacts from eye movements, blinks, heartbeats, and muscle movements. A manual quality control step follows the automated classification.

After the preprocessing, the EEG recordings were post-processed and assessed for comparability [13]. The proportion of time removed from data due to artifact or technical issues (extreme voltage variance, low correlation with neighboring channels, artifact identified by ICA decomposition) was similar across sites. The average channel retention (range: 77%-82%), the spatial variance in the retained and rejected independent components, and the power spectrum profile were also similar across sites.

### 2.3. Annotations of segments with eye-movement artifacts and clean EEG

#### 2.3.1. Eye-movement segments

We used an interactive plot generated with MNE-Python (version 1.3.1) [23, 24] to manually annotate the EEG recordings for clean and eye-movement segments (Fig. 1). We selected segments with horizontal and vertical eyemovement artifacts (orange windows in Fig. 1) based on previous experience and reference images from prior studies [25]. These annotations were made using the channels that are most susceptible to eye-movement artifacts in the EGI 128-channel Hydrocel Geodesic net (E1, E8, E14, E21, E32, E125, E126, E127, and E128). Video recordings or direct measurements of eye movements (i.e., eye-tracking) were not available. To verify the accuracy of the annotated eye-movement windows, we plotted scalp topographic plots of amplitude. The amplitude for ocular artifacts is expected to be higher in the frontal regions [26], with a left-to-right polarity inversion in the case of lateral saccades. The final selection was based on both temporal and topological profiles.

**Figure 1:**
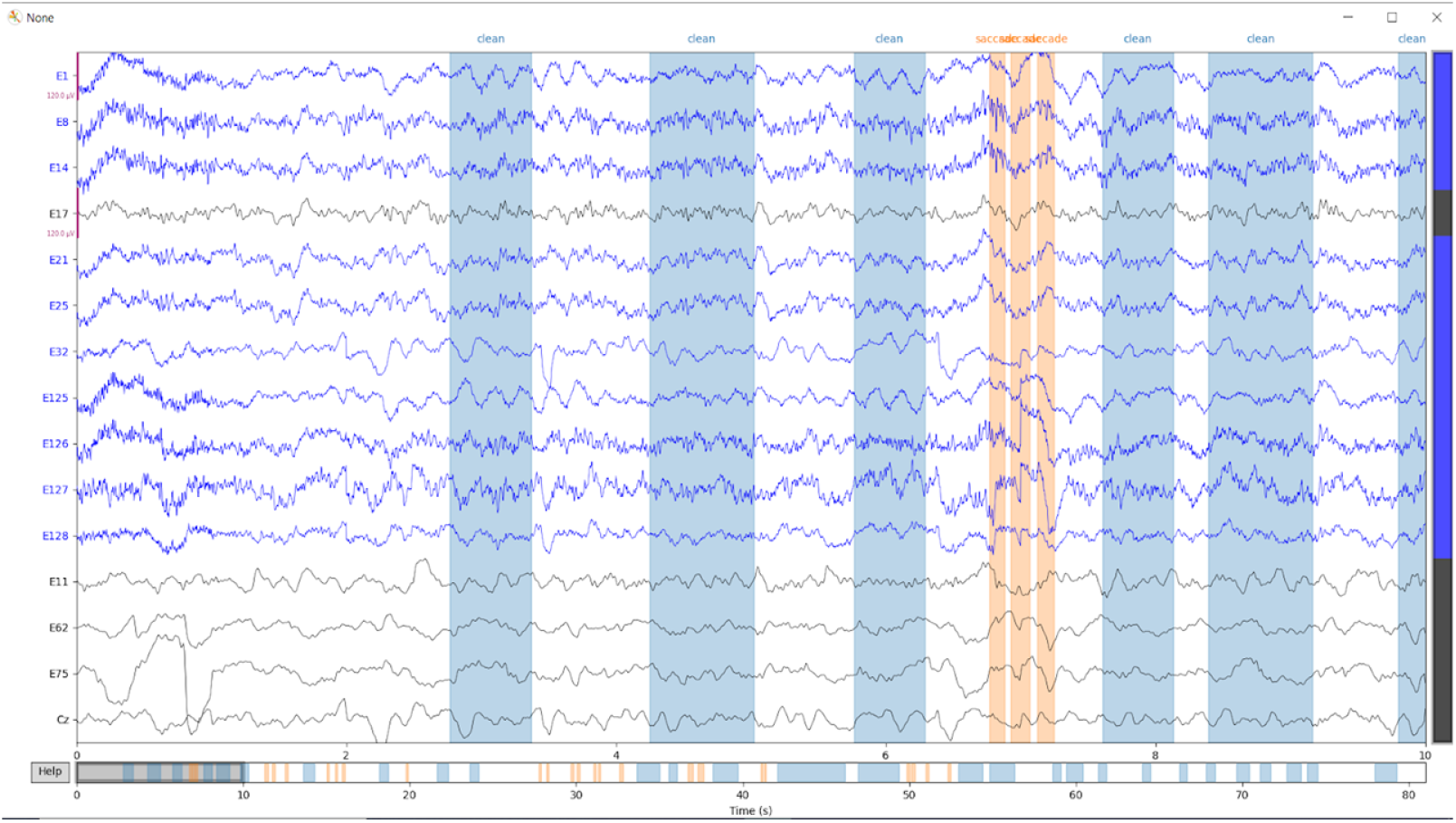
Interactive window used to annotate eye-movement segments (light orange sections) and clean segments (blue sections). Channels used to annotate eye movements are indicated by the blue lines, while clean segment reference channels are indicated by the black lines.

#### 2.3.2. Clean segments

We marked segments without discernible artifacts as clean. For this, we visually inspected channels Fz (E11), Cz (REF), Pz (E62) and Oz (E75). Before marking a segment as clean, we cross-checked patterns of artifacts with an adult EEG atlas of independent components corresponding to various artifact types and brain activity [27]. Segments without discernible artifacts were marked as clean (blue windows in Fig. 1). In a prior study, the authors reviewed EEG studies in infants and children, and recommended a minimum of 30s of clean segments for evaluating artifacts in pediatric EEG [26]. We followed this recommendation and included a minimum of 30s of clean segments per recording.

### 2.4. ICA-based artifact correction

ICA can be used to correct artifacts but depends on three assumptions: i) Signals (S) captured by the electrodes reflect multiple linearly mixed and statistically independent components (sources) of brain activity and artifacts; ii) the number of channels is greater than the number of components; iii) there are negligible volume conduction delays [28]. The linear model for S obtained through ICA is given as:

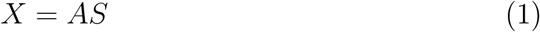

where X stands for the linear mixture of signals *s*_*i*_ in *S* = (*s*_1_, *s*_2_, *s*_3_, …, *s*_*n*_), and A is the unknown mixing matrix. S is approximated by finding the unmixing matrix *A*^*−*1^ (see [22] for details):

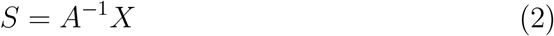

ICA was computed and visually validated before this study as part of the EEG-IP project [13, 5]. All independent components (IC) indicative of artifacts (not specific to eye movements) were selected for rejection. These components included artifacts from the eyes (movements and blinks), heart, muscles, line noise, and channel noise, as defined in ICLabels. Components were automatically labeled for rejection if they were not labeled as “brain” or “other” with at least a 0.3 probability. This automated labeling was manually reviewed and corrected during a quality control step. The set of components labeled for rejection was removed after selecting segments with eye movements or clean EEG but before assessing the performance of ICA.

### 2.5. AB-based artifact correction

The AB algorithm relies on a threshold *θ*_*opt*_ to distinguish between brain signals and high amplitude artifacts. Since this threshold likely depends on recordings and subjects (e.g., due to differences in electrode impedance or head tissue conductivities causing systematic biases in EEG amplitude), we developed a procedure to calculate the optimal threshold *θ*_*opt*_. We define this threshold as the voltage at which artifacts due to eye movements are maximally removed while minimizing distortion to clean segments. Before starting the calculation, we examined clean and eye-movement segments for each recording and chose only channels with sufficient data (i.e., more than 30 seconds of clean EEG) for analysis. For each channel, the following procedure was followed:

1. The maximum amplitude in each annotated segment (eye-movement and clean) was calculated. These maximum amplitudes were used to construct a range of amplitudes as thresholds to be tested. The range consisted of all thresholds, in increments of 1 *μ*V, from the lowest to the highest maximum amplitude recorded across channels.
2. For each threshold, the Mean Absolute Correction (MAC) was calculated for each annotation as follows:

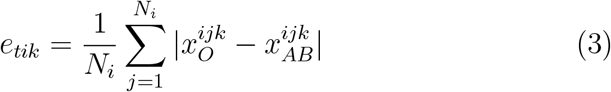

where *e*_*tik*_ is the total error, *t* is the threshold index, *i* denotes the index of the annotation, *k* identifies the channel, *N*_*i*_ stands for the number of time samples in annotation *i, j* is the time sample index, and *x*_*O*_ and *x*_*AB*_ represent the amplitude of the original and the AB-processed signal, respectively.
3. The average MAC was calculated across clean and eye-movement segments, respectively.
4. The average MAC for each type of segment was normalized to be on the same scale (Fig. 2A):

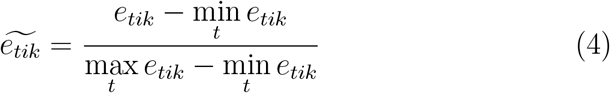
5. The total error (*E*_*tk*_) at each threshold takes into account the average MAC (*e*_*tik*_) for segments with and without artifacts and is defined as follows

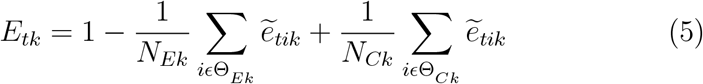

where Θ_*Ek*_ and Θ_*Ck*_ stand for the set of segments with eye movements and clean EEG, respectively, and *N*_*Ek*_ and *N*_*Ck*_ stand for the number of segments in these two sets. Note that a large “error” for eye-movement segments is desirable (i.e., the cleaned signals should be different than the raw signals) whereas a small error is desirable for the clean segments (i.e., the artifact rejection should not distort the signals in such epochs), hence the subtraction of the former and the addition of the latter type of errors. Also, note that an equal weight for the contribution of both types of errors is implicitly used by not further weighting these two types of errors.
6. The optimal threshold for the channel *k* corresponds to the minimum of all total errors calculated (Fig.2B):

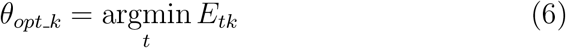
7. Since the original algorithm uses a common threshold for all channels, the optimal threshold for the recording was calculated as the average optimal threshold of all selected channels using

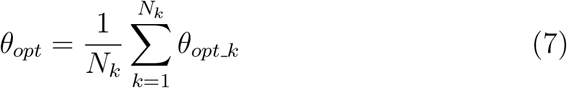

where *N*_*k*_ stands for the number of channels used for threshold evaluation.

Given a two-dimensional EEG matrix *X* of sizes *n × T*, with *n* being the number of channels and *T* being the number of time points, a matrix *Y* is constructed from *X* by zeroing values exceeding *θ*_*opt*_ [29, 30]. Then, a blocking matrix *B*_*opt*_ is constructed as follows:

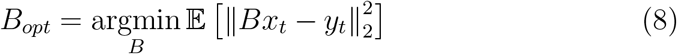

where *x*_*t*_ and *y*_*t*_ refer to the *t*^*th*^ column of matrices *X* and *Y*, respectively, and 𝔼 stands for the expectation operator. Additional details on the calculation of *B*_*opt*_ are available in [31].

**Figure 2:**
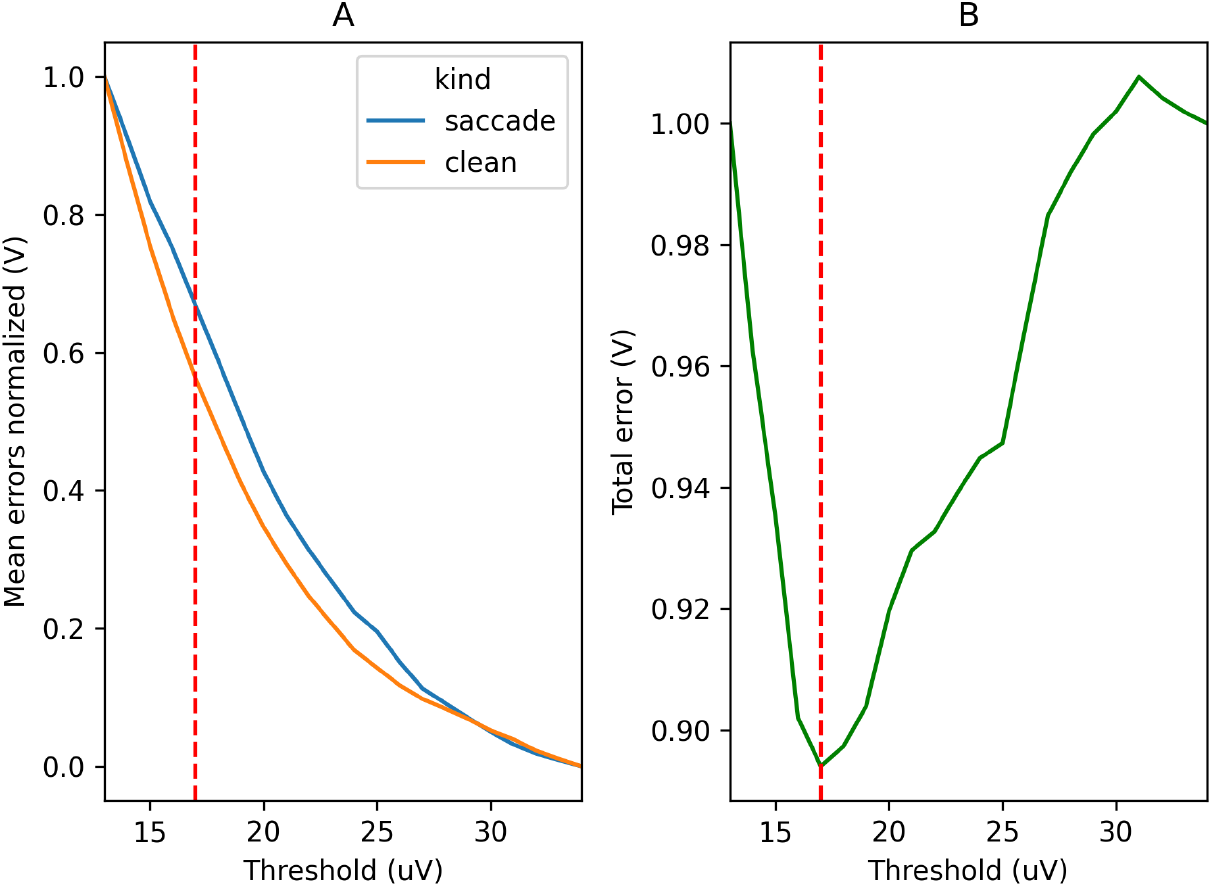
A) Normalized MAC across all tested thresholds for channel E1 in a typical recording. B) The optimal threshold for this channel is 17 *μ*V, i.e., the threshold corre-sponding to the minimum of all total errors calculated for this channel, *θ*_*opt k*_.

**Figure 3:**
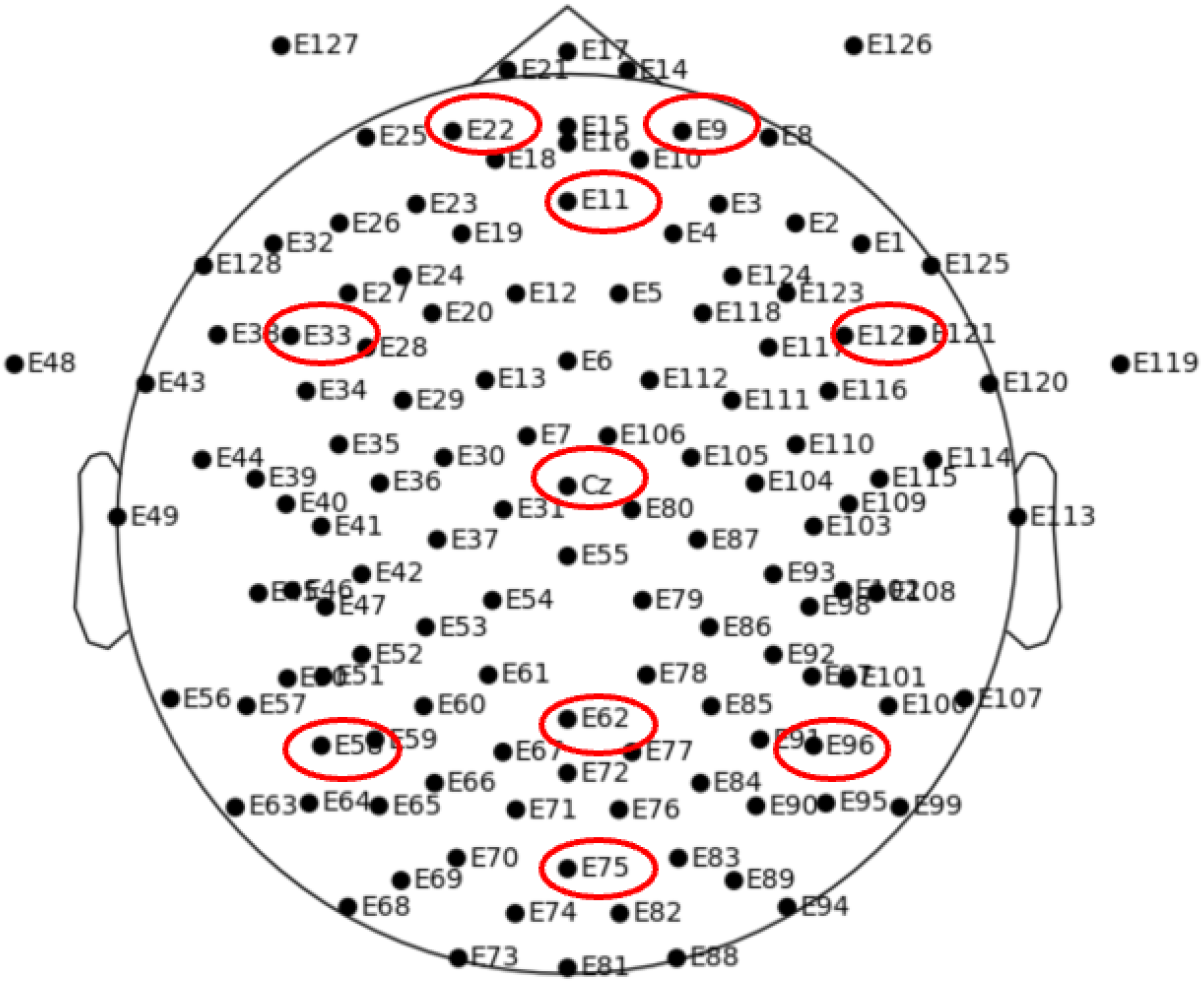
EGI 128-channel Hydrocel sensors net. Sensors circled in red were used for multiscale entropy analyses.

The blocking matrix *B*_*opt*_ utilizes information from the reference matrix *Y* indicating the EEG samples exceeding *θ*_*opt*_. It ‘blocks’ these samples while preserving the samples not exceeding *θ*_*opt*_.

We used the MATLAB script for AB shared by the original authors [12]. We exported to Python the results from MATLAB for comparison with ICA. Portions of code written in MATLAB were integrated into the Python pipeline by running them from Python using the MATLAB engine.

### 2.6. Performance metrics

#### 2.6.1. Proportion of segments effectively corrected by AB and ICA

For each epoch with eye movement, we selected the channel with the most visible artifacts and categorized the application of AB and ICA as either effective or insufficient (i.e., under-correction). The rater was blind to the signal origin (i.e., AB or ICA). To make this annotation procedure systematic and efficient, we implemented a custom interactive window displaying the corresponding time series and accepting sequential annotations through keystrokes. The following procedure was used:

1. For each annotated eye-movement segment, the time series of the original signal was overlaid to the signals corrected with ICA and AB.
2. For each segment, AB- and ICA-corrected traces were randomly assigned a color (red or blue).
3. The red and blue plots were rated as effectively or insufficiently corrected for an eye-movement artifact. The authors (DS and COR) discussed various examples and agreed on criteria for effective correction vs under-correction. An epoch was rated as under-corrected if an eye-movement artifact was still discernible on the plot after correction.
4. At the end of this procedure, blinded scores were automatically associated with their true labels (i.e., ICA or AB) based on a log of the random draws. The proportions of segments where AB and ICA effectively or insufficiently corrected for eye movements were calculated.

We did not manually annotate clean segments as the ideal outcome is known in this case (i.e., it is identical to the original signal since these segments are considered free from artifacts). Any deviation from the original signal is considered an over-correction, and this deviation can be measured quantitatively, e.g., through the SNR. We hypothesized that AB and ICA would correct equally effectively for eye-movement artifacts, i.e., *Pe*_*AB*_ *≈ Pe*_*ICA*_, where *Pe* stands for the proportion of effectively corrected eye-movement segments.

#### 2.6.2. Signal-to-noise ratio (SNR)

The SNR measures the relative amount of noise present in a signal. In our calculations, the signal refers to the processed EEG after applying a cleaning algorithm (i.e., AB or ICA) while noise refers to the portion of EEG removed during preprocessing. This definition implies that large SNRs are desirable for clean segments, while small SNRs are desirable for segments with artifacts. We calculated SNR for each segment by first calculating the root-mean-square (RMS) for the segment processed with AB or ICA, using (9) with *y*_*i*_ being the processed segment (signal) and *n* the number of samples in the segment. Then, we calculated the RMS for the noise using (10) with *x*_*i*_ being the original, unprocessed segment.

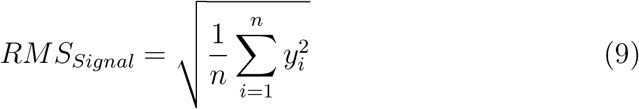

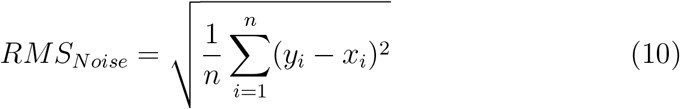

Finally, the SNR was calculated as the quotient of the two RMS values, using a base-10 logarithmic scale and multiplying by 10 to express this ratio in decibels (dB), as is usual:

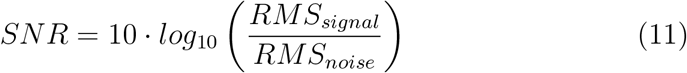

SNR calculations were implemented in Python. An SNR of 0 dB indicates signal and noise RMS values of equal amplitude, whereas a value of 10 *·* 2 = 20 dB indicates a signal 10^2^ = 100 times as large as the noise. Since we defined the noise as what is removed from the raw data by the artifact correction algorithm, a large “noise” (hence, a low SNR) is desirable when assessing the performance of the algorithm on epochs with artifacts, while this “noise” would ideally be zero (hence, a large, theoretically infinite, SNR) for epochs without artifacts, indicating that the algorithm did not distort clean signals. We hypothesized that AB and ICA would correct equally effectively for eye-movement artifacts (*SNR*_*AB*_ *≈ SNR*_*ICA*_), but AB would cause less distortion to the clean segments compared to ICA (*SNR*_*AB*_ *> SNR*_*ICA*_).

#### 2.6.3. Power spectral density (PSD)

The PSD measures the energy contained in a signal at different frequencies. We calculated this value from averaged-reference EEG for all segments, channels, and recordings. For each recording, we cropped individual segments and concatenated them into separate EEG matrices for clean and eye-movement segments. We used the *psd welch* function in MNE-Python [24] to estimate the spectrum with the Welch method. In short, a Hamming window was applied on the segmented windows with 50% overlap. Then, for each channel, we averaged the periodograms calculated for every window. PSD values were *log*_10_-transformed to account for skewed distributions. For our analyses, we computed the total PSD power for the following frequency bands: delta (2-4 Hz), theta (4-6 Hz), low alpha (6-9 Hz), high alpha (9-13 Hz), beta (13-30 Hz), and gamma (30-48 Hz) [32].

We hypothesize that AB and ICA will correct equally effectively for eye-movement artifacts and, consequently, that we will observe an equivalent change in PSD for both algorithms with respect to the original signal (i.e., |*PSD*_*ICA*_*−PSD*_*original*_| *≈* |*PSD*_*AB*_ *−PSD*_*original*_|). We also hypothesize that AB will cause less distortion to the clean segments compared to ICA, which will be visible as a lower change in PSD (i.e., |*PSD*_*ICA*_ *− PSD*_*original*_| *>* |*PSD*_*AB*_ *− PSD*_*original*_|).

#### 2.6.4. Multiscale Entropy (MSE)

MSE is a non-linear measure of signal complexity. We calculated MSE using the *entropy multiscale* function from the Python neurokit2 package [33]. The calculation coarse grains the EEG signal for multiple scales [34] using

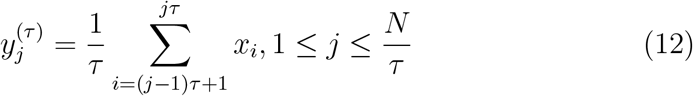

where *y* = *{y*_1_, …, *y*_*j*_, …, *y*_*N*_ *}* is the coarse-grained time series, *τ* is the scale factor, and *x* = *{x*_1_, …, *x*_*i*_, …, *x*_*N*_ *}* is the original signal. We calculated the number of scales per signal using the following formula:

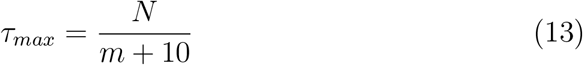

where *N* is the length of the signal. The *m* coefficient is called the embedding dimension, and it refers to the number of consecutive time points accounted for in the equation of sample entropy. An examination of entropy using cardiac and respiratory data found that an *m* = 2 or 3 yielded optimal results [35]. Using *m* = 2, we calculated the sample entropy [35, 36] for the coarse-grained signal at each scale using the following equation:

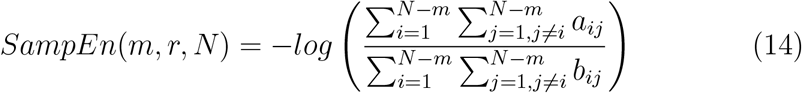

where

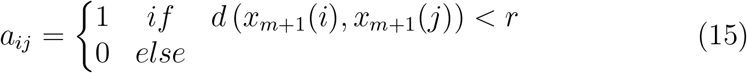

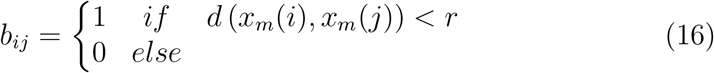

with *d*(*x, y*) = max_*i*_(|*x*_*i*_ *− y*_*i*_|) is the Chebyshev distance between time series *x* and *y*, and *x*_*m*_(*i*) is a segment of the signal *x* defined as the time series *{x*_*i*_, *x*_*i*+1_, *x*_*i*+1_, …, *x*_*i*+*m−*1_*}*. The factor *r* is known as the tolerance, and it limits the distance between the two points *x* and *y*. Based on tests with heart rate and neural data, an *r* value between 0.1 and 0.25 times the standard deviation of the data is recommended. We calculated MSE as the sum of the sample entropy across all scales and focused on its value in ten channels well-distributed across the scalp (Fig.3). We initially calculated entropy using a *τ* that varied by recording based on length. We noticed that differences in entropy between processed and original recordings reached a plateau after 20 scales for clean segments. Hence, we plotted differences in entropy for 20 scales across both clean and eye movement segments, for comparability.

As for the other metrics, we hypothesize that an equivalent efficiency of AB and ICA for artifact correction will be reflected in similar changes in MSE for eye-movement segments (i.e., |*MSE*_*AB*_ *−MSE*_*original*_| *≈ MSE*_*ICA*_ *− MSE*_*original*_|), but that a lesser distortion from AB will result in |*MSE*_*AB*_ *− MSE*_*original*_| *<* |*MSE*_*ICA*_ *− MSE*_*original*_| for clean segments.

### 2.7. Statistical analyses

To compare the difference in proportions of under-corrected and effectively corrected artifacts for each algorithm, we performed a McNemar’s test for difference in proportions. We used the non-parametric Wilcoxon signedrank test to determine the statistical significance of differences in means between artifact correction methods for SNR, PSD, and MSE. Statistical difference between the methods was assessed separately for eye movement segments and clean segments. A cluster-based permutation test using the physical layout of the EEG electrodes to build an adjacency matrix was applied to correct for multiple comparisons [37]. The adjacency matrix was built using Delaunay triangulations. This method uses the 2D electrode locations to construct triangular meshes by maximizing each triangle’s minimum angle. Clusters are identified by identifying sensor locations that fall on the circular area around triangles, once the mesh is constructed [38]. We set a cluster forming threshold at t=434, corresponding to an alpha=0.05 and n=50 for the Wilcoxon signed rank distribution [39]. We computed 1000 random permutations of data between signals processed with AB and signals processed with ICA. We defined clusters of SNR values as groups of tests exceeding the significance threshold in the channel space, yielding 129 comparisons. For PSD, we defined clusters in the channel and frequency band space (129 × 6 comparisons), while we defined them in the channel and scale space for MSE (129 × 20 comparisons). We used the Python libraries StatsModels and SciPy [40, 41] for statistical analyses and MNE-Python [24] for cluster-permutation tests.

## 3. Results

We compared eye-movement artifact removal and clean segment preservation by AB and ICA based on the proportion of eye-movement segments effectively corrected, SNR, PSD and MSE.

### 3.1. Proportion of eye-movement segments effectively corrected by AB and ICA

We performed a blinded comparison of the level of correction achieved by AB and ICA for eye-movement segments. To compare the difference in proportions of under-corrected and effectively corrected artifacts by each algorithm, we performed a two-tailed McNemar’s test for difference in proportions. In segments with eye movements, there was a lower proportion of instances where AB (0.34 *±* 0.02) effectively corrected for ocular artifacts compared to ICA (0.54 *±* 0.02). This difference was statistically significant (*χ*^2^ = 32.52, *df* = 1, *p* = 1.182 *×* 10^*−*8^). These findings suggest that ICA corrects eye-movement artifacts better than AB, not supporting our prediction of equal effectiveness of AB and ICA for artifact correction.

### 3.2. SNR

We calculated the SNR in clean and eye-movement segments to assess the efficacy of ICA and AB in the time domain. In both cases, the SNR increased more for AB than ICA (Fig. 4). The difference is most pronounced in the left frontal, central, and right-posterior regions. To compare the SNR following AB vs ICA processing, we performed a two-tailed Wilcoxon signed-rank test followed by a cluster-based permutation test, separately for the eye movement and clean segments. We found one significant cluster including all 128 channels (*p <* 0.001, permutation test, family-wise error controlled) for both segment types. These trends suggest that ICA corrects better than AB for eye movements (i.e., stronger artifact suppression in ICA increases the “noise” denominator, reducing the SNR). However, the higher SNR for AB in clean segments indicates that this algorithm causes less distortion for EEG compared to ICA (i.e., less artifact removal in artifact-free segments is associated with less signal distortion and higher SNR). Our findings do not support our first prediction that the SNR following AB and ICA would be similar in the eye-movement segments. However, they support our second prediction that the SNR would be greater after AB than ICA in the clean segments.

**Figure 4:**
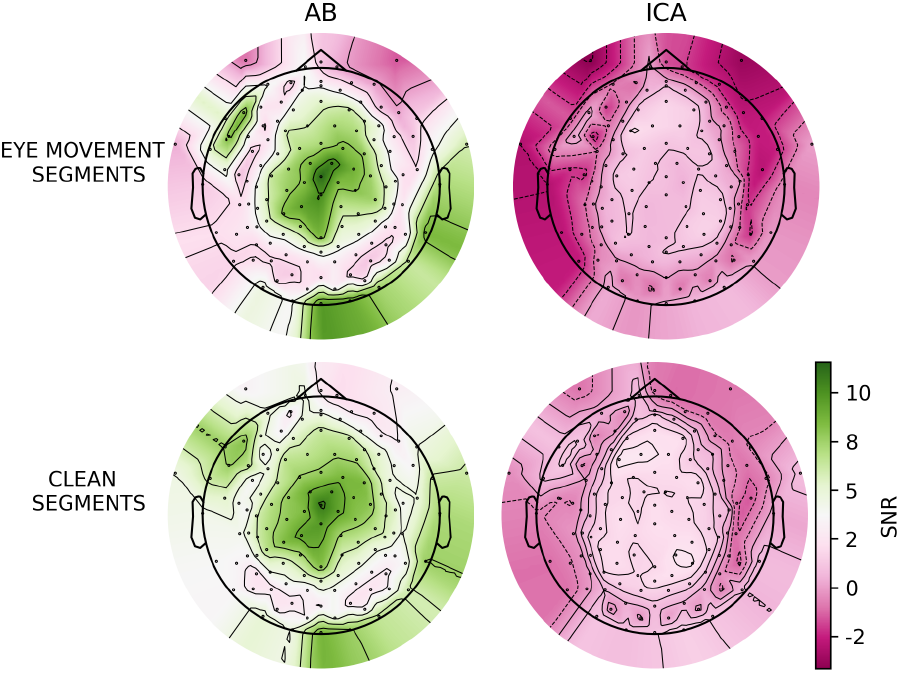
Scalp topographic plots of average SNR following artifact correction using AB and ICA, for segments with eye movements and clean EEG.

### 3.3. PSD

To assess the efficacy of AB and ICA in the frequency domain, we compared the change in PSD associated with AB and ICA. We calculated log_10_PSD in the eye-movement and clean segments for the original, AB-corrected, and ICA-corrected signals across different frequency bands between 2 and 48Hz. For each recording, clean and eye-movement segments were concatenated before calculating log_10_PSD. The average length of concatenated segments was 3.29s for eye movements and 27.37s for clean EEG.

Fig. 5 shows the difference in log_10_PSD between the original and the corrected signals for both segment types. In the eye-movement segments, except for the gamma band, a qualitatively similar change in PSD was observed across all frequency bands, in keeping with our hypothesis that both methods would perform equally. However, a cluster-based permutation test between *log*_10_|*PSD*_*ICA*_ *− PSD*_*original*_| and *log*_10_|*PSD*_*AB*_ *− PSD*_*original*_| revealed a significant cluster (*p <* 0.001, permutation tests, family-wise error controlled) containing 86 electrodes distributed across the scalp and all six frequency bands (see Fig. A2).

**Figure 5:**
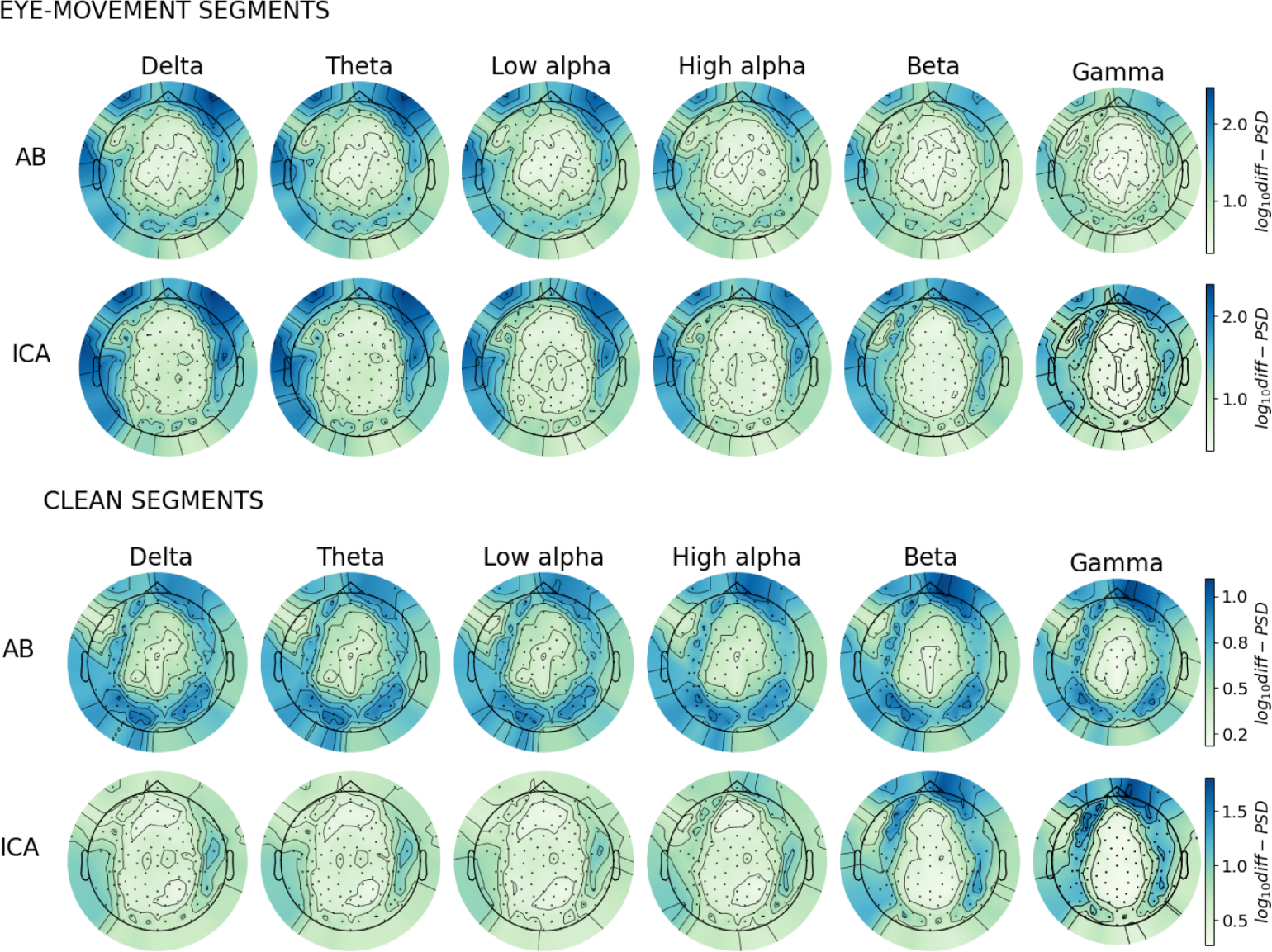
Scalp topographic plots depicting the differences in log_10_PSD between processed and original signals for both segment types. log_10_PSD values have been calculated by averaging across 50 participants for each frequency band and channel. See Fig. A2 for the results from the statistical tests (cluster-based permutation) between AB and ICA.

For clean segments, *log*_10_|*PSD*_*ICA*_ *− PSD*_*original*_| in the beta and gamma bands is greater than *log*_10_|*PSD*_*AB*_ *− PSD*_*original*_|. This observation suggests that ICA distorts more clean segments than AB, as per our prediction. The corresponding cluster-based permutation test revealed a single significant cluster (*p <* 0.001, permutation tests, family-wise error controlled) containing 107 electrodes across the scalp and all six frequency bands (see Fig. A2).

### 3.4. MSE

In eye-movement segments, |*MSE*_*ICA*_ *− MSE*_*original*_| and |*MSE*_*AB*_ *− MSE*_*original*_| are similar in channels F8, Fp1, Oz, and T6. There are variations by scale for the other channels (Fig. 6). These findings partially support our prediction that entropy would change by a similar amount following correction by both algorithms. We used a cluster-based permutation to compare between |*MSE*_*ICA*_ *− MSE*_*original*_| and |*MSE*_*AB*_ *− MSE*_*original*_| and found a single significant cluster (*p <* 0.001, permutation tests, family-wise error controlled) containing 32 electrodes distributed across the frontal and central regions and all 8 scales.

**Figure 6:**
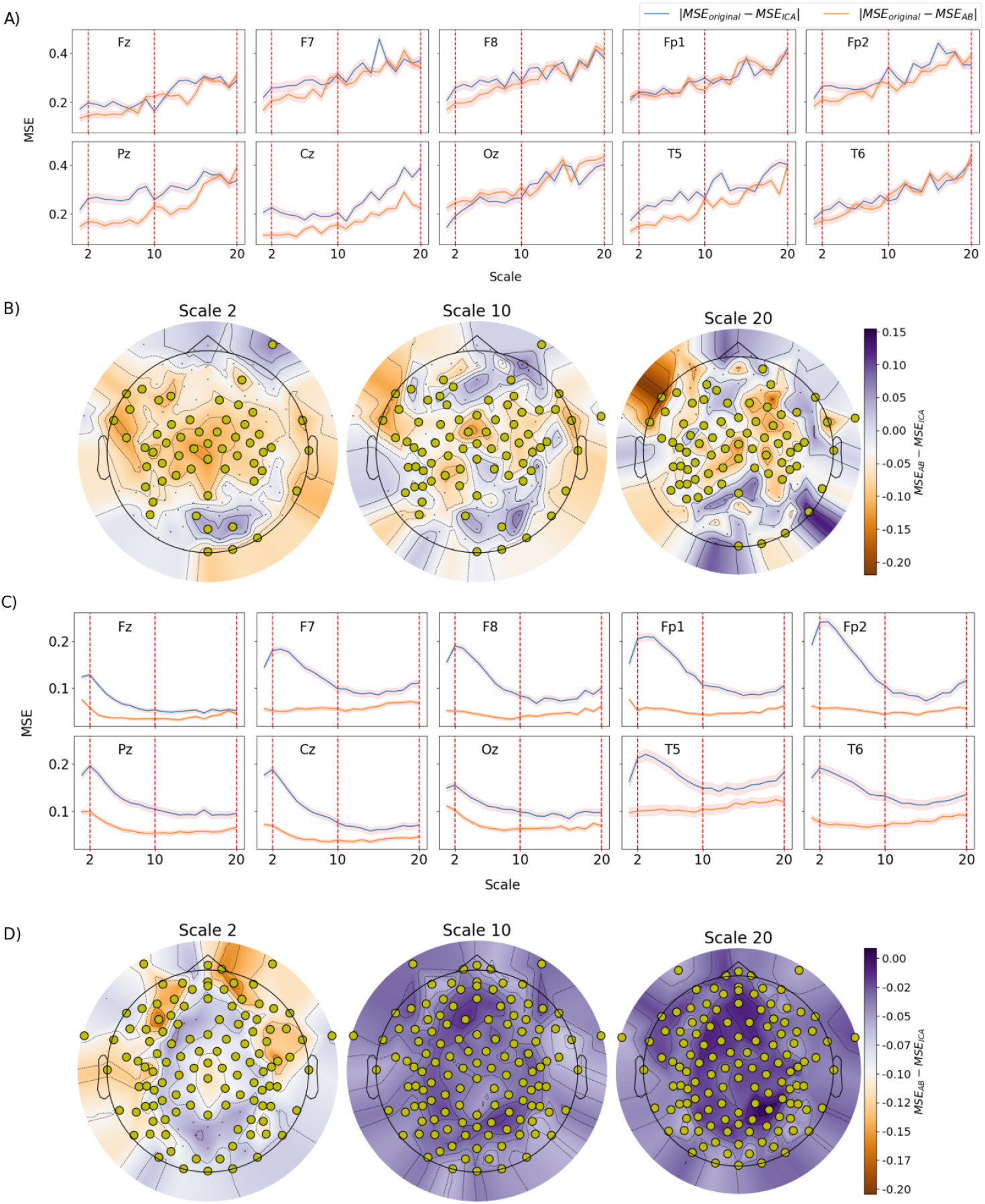
Line-plots show the difference across scales in MSE with respect to original signal following correction by AB vs ICA in A)Eye movement segments, C)Clean segments. Shaded regions in indicate the 95% confidence intervals. The topographic plots show the relative differences in MSE, across channels, at specific scales in B)Eye movement segments, D)Clean segments. MSE values have been averaged across 50 participants and 20 scales for each of 129 channels. Highlighted circles depict the cluster of electrodes with significant differences in MSE between AB and ICA.

In the clean segments, all channels show that |*MSE*_*AB*_ *− MSE*_*original*_| is lower than |*MSE*_*ICA*_ *− MSE*_*original*_|. These findings support our hypothesis that AB would cause less distortion in the clean segments compared to ICA. The corresponding cluster-based permutation test found one significant cluster (*p <* 0.001, permutation tests, family-wise error controlled) containing all 128 electrodes and all 20 scales.

## 4. Discussion

This study applied AB and ICA on a large dataset (N=50) of infant EEG to compare their effectiveness for removing eye-movement artifacts without distorting neural signals. The proportion of effectively corrected segments indicated that ICA corrected eye-movement artifacts better than AB, that is, ICA has better sensitivity to eye-movement artifacts. SNR and PSD suggested that AB and ICA correct for eye-movement artifacts equally effectively (i.e., they are as sensitive), while MSE results were mixed. Our results suggest that AB has better specificity (causes less distortion to clean segments) than ICA, as assessed via SNR, PSD, and MSE.

Our results on the proportion of effectively corrected segments are contrary to a prior qualitative comparison of AB and ICA that found AB to be more effective than ICA for removing eye-movement artifacts in infant EEG [31]. However, since this previous study relied only on one recording, results from that prior study are likely unreliable.

In agreement with the expectation for the eye-movement artifacts to be dominant in the frontal regions, the amount of artifact correction performed by both algorithms depended on the regions of the scalp, with lower SNR (i.e., more artifacts removed) in frontal channels than in central, parietal, and temporal channels. Many studies evaluating ICA and other ocular artifact removal algorithms have calculated the SNR on simulated rather than real EEG to benefit from a known ground truth about the composition of the signal and noise. While a comparison with real EEG is essential because of its ecological validity, simulations provide complementary evidence by characterizing how well the algorithm performs according to the ground truth about the (simulated) brain activity and eye activity [42]. A potential next step to strengthen our analyses is to compare AB and ICA using simulated EEG generated using ocular artifacts extracted from a real dataset. Simulated recordings could be generated by extracting segments of eye artifacts from EEG recordings [43].

Due to differences in definitions of SNR, values from prior studies are not directly comparable with those reported here [44]. We also believe this study to be the first to calculate the difference in MSE between the original and corrected versions of eye-movement artifact segments.

PSD is one of the metrics most commonly evaluated in studies assessing the performances of artifact removal algorithms on real EEG. We found that ICA distorts more clean segments in the beta and gamma frequency bands than AB, as per our prediction. The reduction in power at these higher frequencies may be related to components associated with other sources of noise, such as muscle movement [45]. Unfortunately, the manual rejection of independent components for cleaning the signals was done before this work and was not specific to EOG artifacts. These also included artifacts from the heart, muscles, line noise, and channel noise, as defined in ICLabels.

This study aims to provide critical information to support researchers in deciding between ICA and AB for artifact removal. Using these four metrics, we have provided a comprehensive comparative analysis of the performance of these two approaches in terms of their effectiveness at removing eye-movement artifacts (sensitivity) and their ability not to distort clean signals (specificity).

Our study is limited by the absence of an independent measure of eye movements. Hence, manual annotations of segments with eye movements were performed based on templates of the characteristic appearance of these artifacts. The visual inspection used to calculate the proportion of effectively corrected eye-movement segments was discussed and validated between the authors (DS and COR), and we iteratively improved the annotation process, until a consensus was reached. However, final annotations were performed by only one rater, which limits their reliability. Besides using multiple raters and computing an inter-rater agreement for the annotations, a future study could use concurrent EEG and eye-tracking to provide a more objective classification.

The optimal threshold calculation for AB was suggested as a way to maximize artifact correction while not distorting clean segments. However, this requires manual annotations of clean and artifactual segments to be done beforehand, which is a drawback. Future studies may want to consider investigating alternative approaches. Such an approach could, for example, use the threshold adopted in the standard AB approach to identify periods with heavy artifacts and with little to no artifacts, use these periods to assess the optimal threshold, and repeat until convergence.

## 5. Conclusion

Our results suggest that ICA might correct better for eye-movement artifacts than AB, while AB causes less distortion to clean EEG than ICA in infant EEG. Our analysis yielded some new insights on methods for benchmarking artifact removal algorithms. To identify more definitively what approach performs best overall for infant EEG, future work should evaluate AB and ICA using infant EEG with eye-tracking. Further, since we found ICA to be more sensitive (better reject artifacts) and AB to be more specific (distort less non-artifact signals) to eye-movement artifacts, automating the assessment of artifact rejection performance in terms of these two metrics (i.e., sensitivity and specificity) and computing these performances across a range of decision threshold could allow computing receiver-operating characteristics (ROC) curves. Such curves could identify more definitively what approach performs best overall, for example, by comparing the area under the ROC curve for these two approaches.

## 6. CRediT authorship contribution statement

**Diksha Srishyla:** Conceptualization, Methodology, Formal Analysis, Writing – original draft, Writing – review and editing, Visualization **Sara Webb:** Writing – review and editing **Mayada Elsabbagh:** Writing – review and editing, Supervision, Funding acquisition **Christian O’Reilly:** Conceptualization, Methodology, Writing - review and editing, Data Curation, Supervision, Funding acquisition **BASIS team:** Investigation, Resources, Project Administration

## 7. Acknowledgements

We want to thank Drs. Julie Scorah and Tim Smith for reviewing drafts of this work. We want to acknowledge James Desjardins, Scott Huberty, and colleagues at Dr. Elsabbagh’s lab for their feedback on the preliminary results. We would like to thank Professor Joseph Rochford for help with statistical analyses. EEG-IP was supported by funding from Brain Canada and FRQ-S. Christian O’Reilly and Diksha Srishyla are supported through startup funds from the University of South Carolina provided to Dr. O’Reilly. Diksha Srishyla was also supported by the Maysie Macsporran fellowship from the Faculty of Medicine of McGill.

Original data acquisition was funded by NIH P50 HD055782 (Webb) and the UK Medical Research Council (Johnson).

## Supplementary Figures

**Figure A1:**
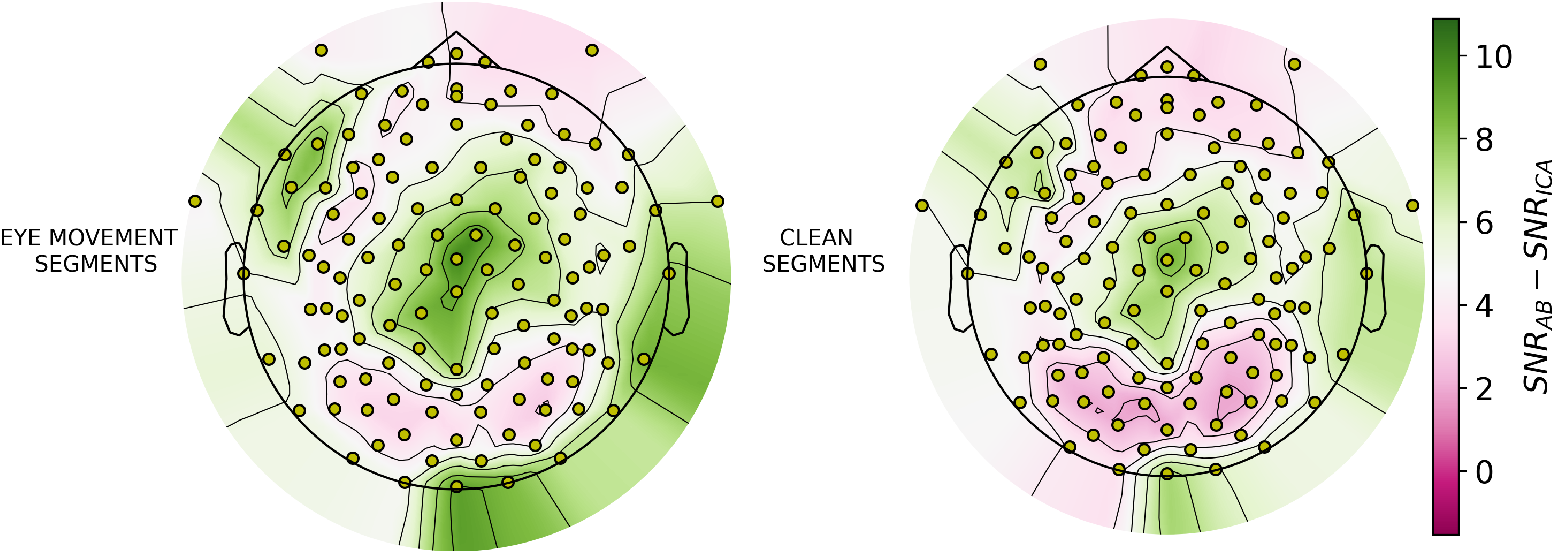
Scalp topographic plots depicting the relative differences in SNR after correction by AB vs after correction by ICA. SNR values have been averaged across 50 participants for each of 129 channels. Highlighted circles depict the cluster of electrodes where there were significant differences in SNR between AB and ICA

**Figure A2:**
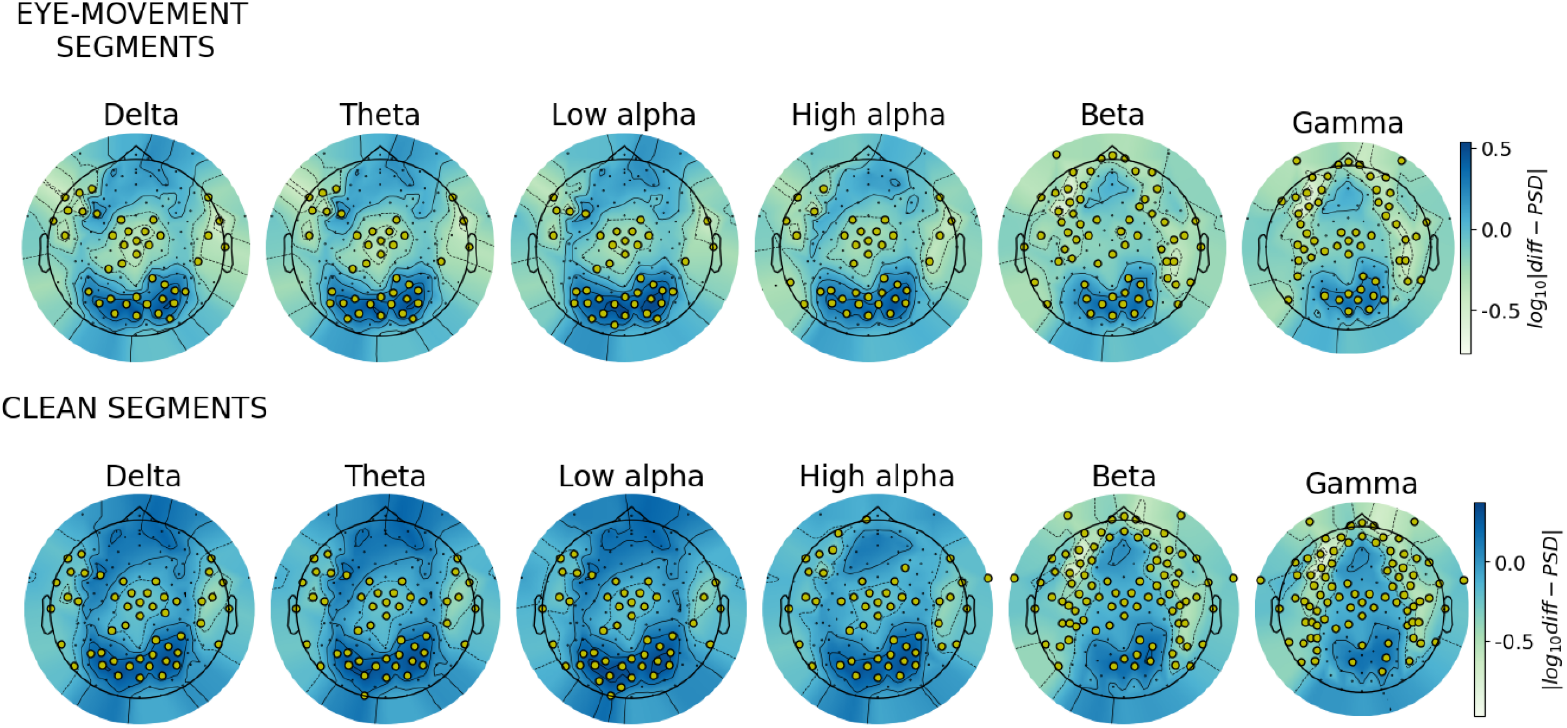
Scalp topographic plots depicting the relative differences in log_10_PSD after correction by AB vs ICA. Relative differences were calculated by first taking the absolute difference between original signal and AB/ICA corrected signal, then calculating the difference between these differences. log_10_PSD values have been calculated by averaging across 50 participants for each of 6 frequency bands and 129 channels. highlighted circles depict the cluster of electrodes with significant differences in log_10_PSD between AB and ICA.

